# Population genomics and the evolution of virulence in the fungal pathogen *Cryptococcus neoformans*

**DOI:** 10.1101/118323

**Authors:** Christopher A. Desjardins, Charles Giamberardino, Sean M. Sykes, Chen-Hsin Yu, Jennifer L. Tenor, Yuan Chen, Timothy Yang, Alexander M. Jones, Sheng Sun, Miriam R. Haverkamp, Joseph Heitman, Anastasia P. Litvintseva, John R. Perfect, Christina A. Cuomo

## Abstract

*Cryptococcus neoformans* is an opportunistic fungal pathogen that causes approximately 625,000 deaths per year from nervous system infections. Here, we leveraged a unique, genetically diverse population of *C. neoformans* from sub-Saharan Africa, commonly isolated from mopane trees, to determine how selective pressures in the environment coincidentally adapted *C. neoformans* for human virulence. Genome sequencing and phylogenetic analysis of 387 isolates, representing the global VNI and African VNB lineages, highlighted a deep, non-recombining split in VNB (herein VNBI and VNBII). VNBII was enriched for clinical samples relative to VNBI, while phenotypic profiling of 183 isolates demonstrated that VNBI isolates were significantly more resistant to oxidative stress and more heavily melanized than VNBII isolates. Lack of melanization in both lineages was associated with loss-of-function mutations in the *BZP4* transcription factor. A genome-wide association study across all VNB isolates revealed sequence differences between clinical and environmental isolates in virulence factors and stress response genes. Inositol transporters and catabolism genes, which process sugars present in plants and the human nervous system, were identified as targets of selection in all three lineages. Further phylogenetic and population genomic analyses revealed extensive loss of genetic diversity in VNBI, suggestive of a history of population bottlenecks, along with unique evolutionary trajectories for mating type loci. These data highlight the complex evolutionary interplay between adaptation to natural environments and opportunistic infections, and that selection on specific pathways may predispose isolates to human virulence.

## Introduction

*Cryptococcus neoformans* is an opportunistic fungal pathogen of enormous clinical importance. An estimated one million new *Cryptococcus* infections occur each year resulting in approximately 625,000 deaths (Park et al. 2009). In Africa, cryptococcosis is the third most common cause of death in patients with HIV/AIDS, and mortality from cryptococcal meningitis exceeds the death rate from tuberculosis (Park et al. 2009). Of the two varieties recognized within the *C. neoformans* species complex, over 95% of cryptococcosis cases are caused by *C. neoformans* var. *grubii* (Chayakulkeeree and Perfect 2006) (serotype A lineage).

Genetic analyses have subdivided *C. neoformans* var. *grubii* into three distinct lineages. The VNI and VNII lineages are highly clonal and globally distributed, while the third lineage, VNB, is genetically diverse yet largely restricted to sub-Saharan Africa (Litvintseva et al. 2006). Environmental reservoirs also differ between these groups; VNB is associated with collection from tree hollows and soil at the base of trees, particularly the mopane tree *(Colophospermum mopane)* which predominates in the African savannah (Litvintseva et al. 2011). VNI is associated with both trees and pigeon excreta (Litvintseva et al. 2011), whereas the environmental reservoir for VNII is not well defined, as the vast majority of known VNII isolates have clinical origin.

The three lineages also differ in the prevalence of each of the two mating types, however, the extent to which this impacts recombination is not well understood. In *Cryptococcus,* the two mating types, a and α, are encoded by a ~100 kb mating type locus (*MAT*) (Lengeler et al. 2002). Sexual reproduction and meiotic recombination can result both from bisexual mating of the opposite mating types as well as from unisexual mating between *MATα* isolates (Lin et al. 2005). VNI and VNII are predominantly of a single mating type (*MATα*) while VNB contains a mix of both mating types (*MATα* and *MAT***a**); the presence of both mating types in VNB has led to the suggestion of more extensive recombination in this group compared to VNI and VNII (Litvintseva et al. 2003, 2006). Previous genetic analyses of *C. neoformans* suggested recombination in both single and multiple mating type populations (Litvintseva et al. 2003, 2006; Hiremath et al. 2008), although these studies were based on AFLP or MLST data and therefore could only provide limited resolution.

*C. neoformans* is an environmental opportunistic pathogen, incapable of human to human transmission or dissemination back to the environment from infected hosts. Therefore direct selection of traits to confer virulence or transmission advantages in humans cannot occur. Rather, traits with increased virulence in humans are the result of coincidental selection (Brown et al. 2012), resulting from interactions of *C. neoformans* with other eukaryotes such as predatory amoeba and nematodes in natural environments (Steenbergen et al. 2001). Isolates collected from the environment have been shown to be less virulent in animal models than those recovered from clinical sources (Fromtling et al. 1989; Litvintseva and Mitchell 2009), suggesting that there are advantageous genotypes for growth in a human host. Understanding the selective forces operating on *C. neoformans* in nature is critical to understanding genetic diversity as it relates to virulence in humans.

Three primary virulence factors for *C. neoformans* include a polysaccharide capsule that inhibits phagocytosis, melanin that protects from environmental stresses, and the ability to grow at human body temperature, 37°C (Bulmer et al. 1967; Kwon-Chung and Rhodes 1986). Numerous additional virulence factors have been identified in *C. neoformans,* such as phospolipase B (Cox et al. 2001), urease (Cox et al. 2000), and a number of signaling cascades (Kozubowski et al. 2009). While the importance of these genes has been confirmed by gene deletion, natural genetic variation encoded within these and other factors could also affect virulence in a quantitative fashion, as genetically similar isolates have been identified that have differential ability to cause disease in animals (Litvintseva and Mitchell 2009).

Previous genomic work has provided reference assemblies of *C. neoformans* (Loftus et al. 2005; Janbon et al. 2014) and the sister species *C. gattii* (D’Souza et al. 2011; Farrer et al. 2015). Here, we leverage the reference assembly of *C. neoformans* strain H99 to conduct large-scale sequencing and variant calling of 387 isolates of *C. neoformans* var. *grubii,* covering the phylogenetic, geographic and ecological breath of the species. We trace the complex evolutionary history of the group using phylogenetic and population genomic analyses, and dissect the genetic differences between clinical and environmental isolates through tests for genome-wide association and selection. For the latter analyses we focus on VNB, as the high genetic diversity and suggested frequent recombination in the lineage, along with ease of isolation from both clinical and environmental sources, make it particularly suited to these tests. Furthermore, we conduct a high-throughput phenotypic screen of 183 VNB isolates to explore how genetic differences relate to responses to environmental stresses. These analyses provide insight into how genetic variation relates to clinical prevalence and how selection in the environment has adapted *C. neoformans* for infection of human hosts.

## Results

### Phylogenetic analysis reveals deep, non-recombining split in VNB lineage

To better understand the population structure and genetic basis of virulence across *C. neoformans* var. *grubii* lineages, we sequenced and identified variants for 387 isolates, including 185 from VNI, 186 from VNB, and 16 from VNII (Methods, **Supplemental Table S1**). A phylogeny estimated from ~ 1 million variable positions showed clear separation of the three monophyletic lineages VNI, VNII, and VNB (**Fig. 1**). However, inspection of the phylogeny also revealed a strongly supported (100% bootstrap support), deep split within the VNB lineage, that matched a division seen in previous analyses of AFLP data (Litvintseva et al. 2003), MLST data (Litvintseva et al. 2006; Chen et al. 2015), four nuclear loci (Ngamskulrungroj et al. 2009), and 23 whole genome sequences (Vanhove et al. 2016). This split was further supported by analysis of population ancestry using STRUCTURE (k=4) and of sequence similarity using principle components analysis (PCA), both of which produced four distinct groups representing VNI, VNII, and two VNB lineages with little to no intermixing (**Fig. 2A,B**). We therefore propose that VNB be subdivided into two distinct lineages, VNBI and VNBII, following the nomenclature of (Chen et al. 2015). VNBI encompassed 122 isolates from this study, including previously studied MATa isolates Bt63 and Bt204, while VNBII included 64 isolates from this study, including previously studied *MAT***a** isolates Bt65 and Bt206 (Litvintseva et al. 2003, 2006; Morrow et al. 2012).

**Figure 1.**
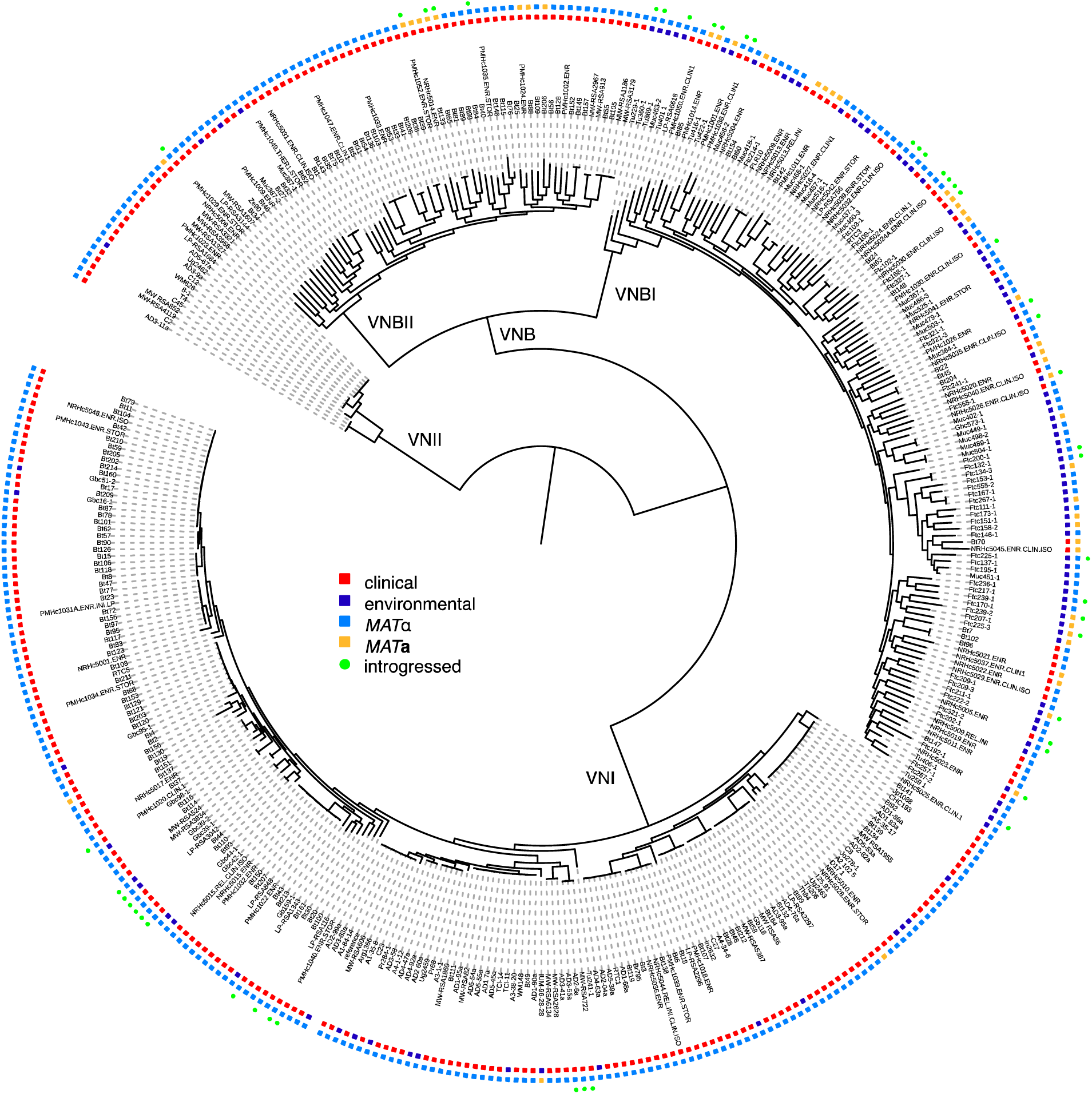
Phylogenomic analysis reveals a deep split within the VNB lineage. We propose that VNB be divided into two lineages: VNBI and VNBII. The phylogeny was estimated from 1,069,080 segregating sites using RAxML (Stamatakis 2014), and the tree was rooted with VNII as the outgroup. All lineages (VNI, VNII, VNBI and VNBII) had 100% bootstrap support. Isolation source (clinical vs. environmental), mating type (*MATα* and *MAT***a**), and presence of >= 50 kb of introgressions from other lineages are also shown. VNBI samples were enriched for environmental isolation sources relative to VNBII (p < 0.0001, Fisher’s exact test).

**Figure 2.**
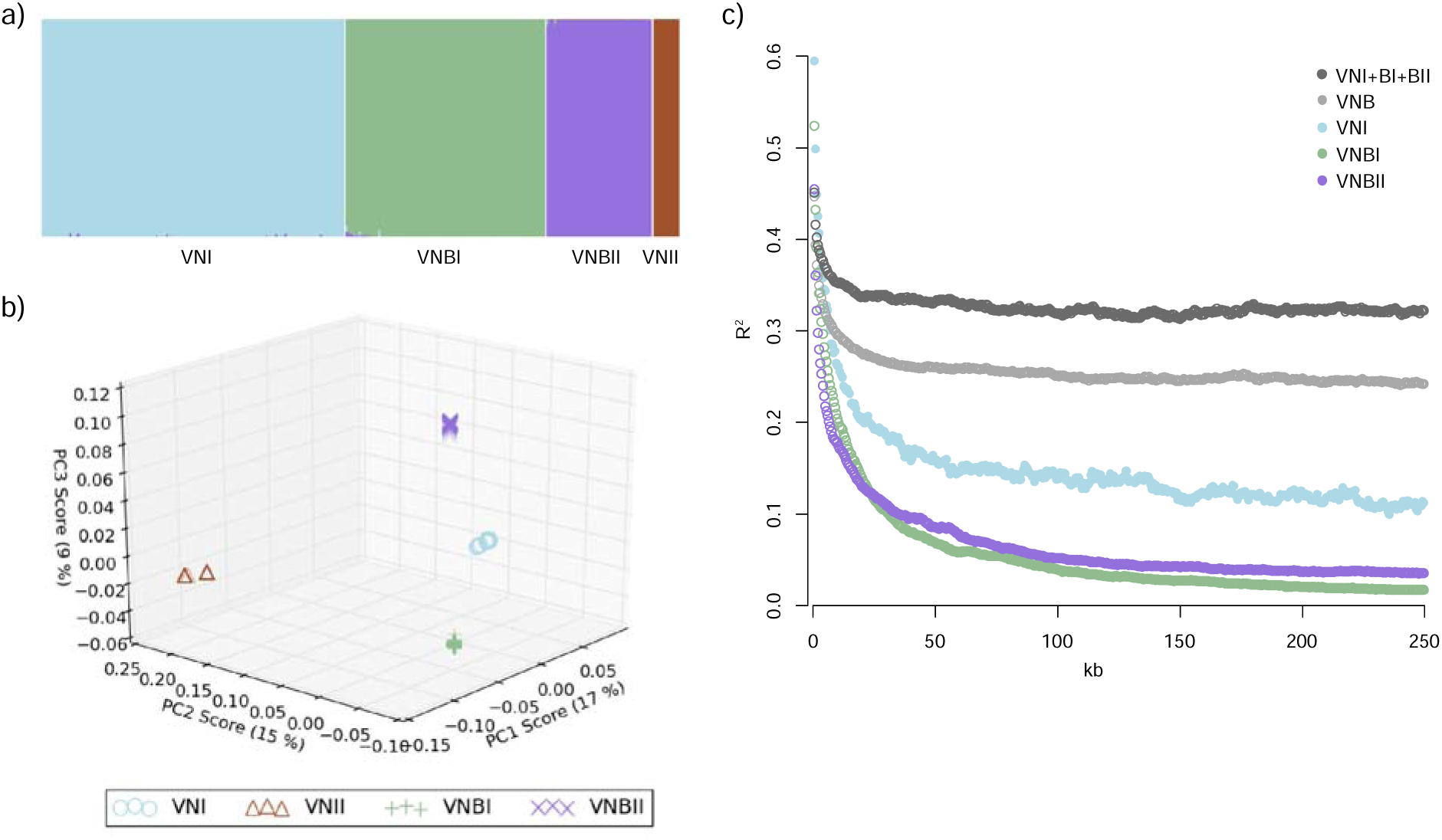
*C. neoformans* var. *grubii* is separated into four distinct non-recombining lineages: VNI, VNII, VNBI and VNBII. A) Results of STRUCTURE analysis (k=4) shows separation of all four lineages; B) Top three principle components from PCA analysis. The third principle component clearly separates VNBI from VNBII; C) Decay of linkage disequilibrium (LD) shows similar rates of recombination within groups and lack of recombination between groups. LD (R^2^) was calculated for all pairs of SNPs separated by 0–250 kb and then averaged for every 500 bp. LD values for each window were then calculated by averaging over all pairwise calculations in the window. The chromosome with the mating type locus was excluded from the calculations.

To determine if these groups represented reproductively isolated, non-recombining lineages, we calculated decay of linkage disequilibrium (LD) for VNI, VNBI, VNBII, all VNB, and VNB combined with VNI. We excluded VNII from the calculations due to small sample size. Neither the combined set of VNB and VNI, nor VNB alone, demonstrated evidence of 50% linkage disequilibrium (LD) decay within the 250 kb range of the analysis (**Fig. 2C**), suggesting limited recombination between members of these populations. However, when split into VNI, VNBI, and VNBII, all lineages demonstrated 50% LD decay in 5 to 7.5 kb (**Fig. 2C, Table 1**). This level of recombination is similar to that previously reported for wild populations of *Saccharomyces cerevisiae* and S. *paradoxus* (Liti et al. 2009).

**Table 1.** Genomic and population characteristics of global sampling of the four lineages of *C. neoformans* var. *grubii.* Listed statistics include sample size, physical distance needed for linkage disequilibrium to decay 50% (LD50), diversity (π), percent of the genome with reduced Tajima’s D (TD), and number and percent of isolates isolated from clinical and environment sources, and of each mating type (*MATα* or *MAT***a**). Given the limited sampling of VNII, many statistics were not determined (ND) or not applicable (NA).

| Statistic | VNI | VNBI | VNBII | VNII |
| --- | --- | --- | --- | --- |
| Sample Size | 185 | 122 | 64 | 16 |
| LD50 (kb) | 7.5 | 6 | 5 | ND |
| $\pi$ | 0.00209 | 0.00360 | 0.00358 | ND |
| % with reduced TD | 1.8 | 12 | 2.4 | ND |
| Metadata |  |  |  |  |
| # Clinical | 156 | 51 | 60 | 13 |
| # Environmental | 24 | 71 | 3 | 1 |
| # *MATα* | 182 | 89 | 55 | 16 |
| # *MATa* | 3 | 33 | 9 | 0 |
| % Clinical/Environmental | NA | 43/57 | 94/6 | NA |
| % *MATa*/$\alpha$ | NA | 27/73 | 14/86 | NA |

In the phylogenetic analysis, VNI was further subdivided into three distinct clades, two of which were globally distributed (VNIa and VNIb) and one of which was restricted to sub-Saharan Africa (VNIc) (**Supplemental Fig. S1**). These three clades were highly supported (100% bootstrap support) in this phylogeny; these subdivisions are consistent with one prior MLST analysis (Litvintseva et al. 2006), but not with two major subdivisions identified in a more recent MLST analysis (Ferreira-Paim et al. 2017) (Methods). The VNIa clade contains the highest diversity (pi=0.00178) compared to VNIc (0.00125) and VNIb (0.00084). STRUCTURE and PCA analyses showed the three clades to be largely distinct (**Supplemental Fig. S1B**), with a small subset of isolates showing evidence of mixed ancestry of all three clades. However, faster LD decay was observed in the full VNI set of isolates compared to that of the individual subclades (**Supplemental Fig. S1C**), suggesting there is active recombination between the VNI subclades and that VNI should be considered a single lineage.

To determine whether there was some level of genetic exchange between the groups below the level of detection of genome-wide LD and STRUCTURE analyses, we repeated the STRUCTURE analysis at 500 kb intervals and identified 46 isolates with evidence of recent introgressions of at least 50 kb from different VN groups (**Fig. 3**, Methods). These introgressions were composed of segments ranging in size from 5 to 260 kb (0.02–1.4% of the genome) and occurred in all pairwise combinations of VNI, VNBI, and VNBII as recipient and donor; VNII donors but not recipients were detected. Thirty-two VNBI isolates contained introgressions, which was significantly more than the 14 VNI isolates and 5 VNBII isolates that contained introgressions (p < 1.32×10^−5^ and p < 0.0033, respectively, Fisher’s exact test). Isolates with introgressions included both mating types and were dispersed across the VNB phylogeny, but within VNI only occurred within the African-specific subclade (Fig. 1). Overall, introgressed regions were not functionally enriched for any specific gene functions; however, the most common introgression in VNBI (present in 14 clinical and environmental isolates, from VNI) encompasses 40 kb and encodes a glycolipid mannosyltransferase involved in capsule synthesis (CNAG_00926) (O’Meara and Alspaugh 2012) and secretome component carboxypeptidase D (Geddes et al. 2015). Conversely, the most common introgression in VNI (present in 8 clinical and environmental isolates, from VNBI) encompasses 60 kb and contains virulence factor *VCX1* (Kmetzsch et al. 2010) and azole drug target *ERG11.* This suggests a low level of mixing of genomic regions with clinically relevant phenotypes may be occurring between lineages in sub-Saharan Africa.

**Figure 3.**
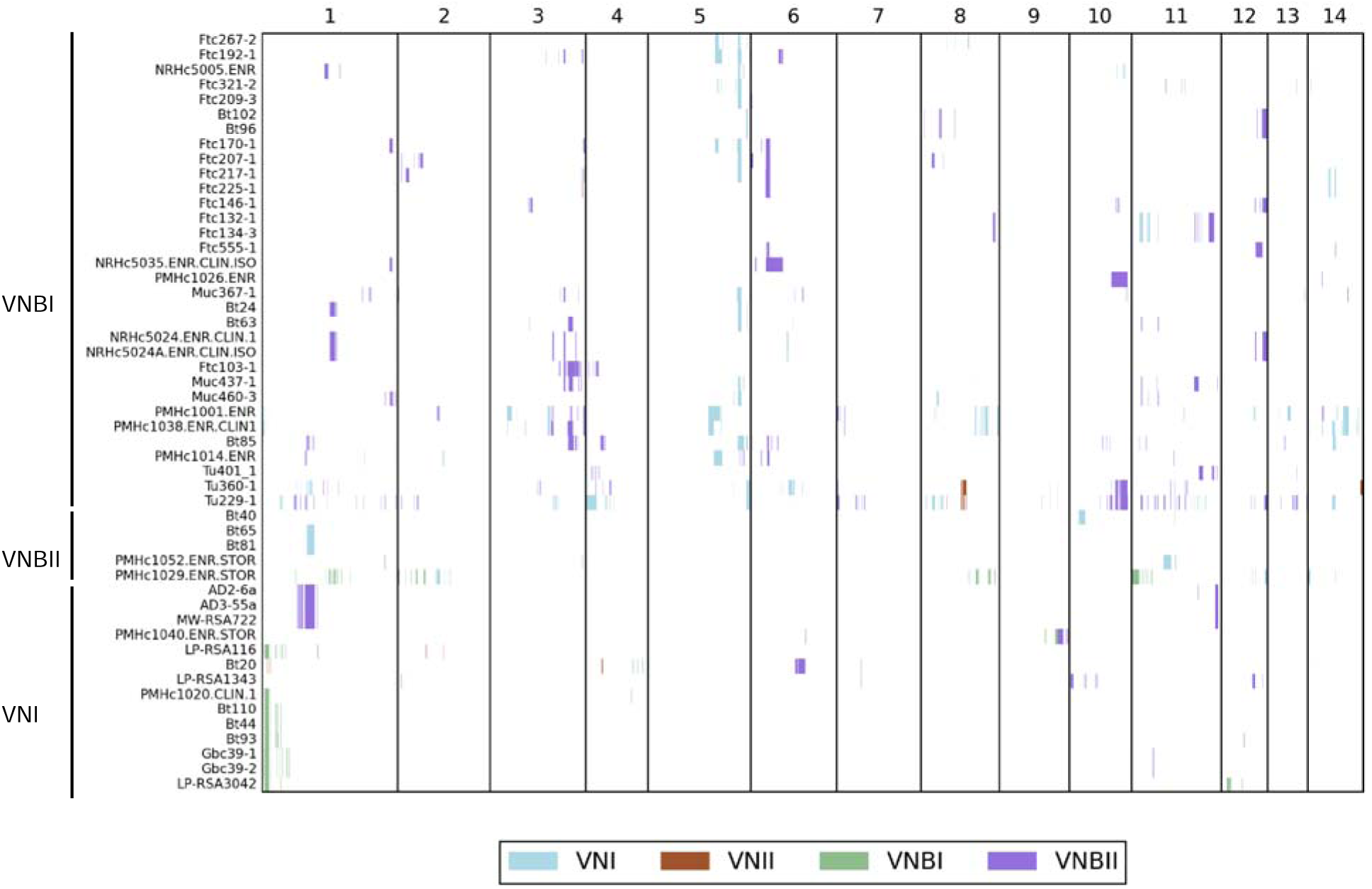
Small introgressions between VNI, VNII, VNBI, and VNBII are dispersed throughout the genome and phylogeny. Introgressions were detected by running STRUCTURE (k=4) on 500 kb blocks, excluding the mating type locus, and the group ancestry of each 5 kb within each block was identified. Recipient genomes are shown on the y-axis while genomic position is shown on the x-axis. White indicates non-introgressed regions while colored blocks indicate introgressed regions. The color key shows which color corresponds to which donor group.

### VNBII isolates are enriched for clinical isolation source relative to VNBI

While the VNB lineage has previously been associated with a high frequency of environmental samples and the **a** mating type, we found differences between the relative frequencies of these properties between VNBI and VNBII. While VNBI contained large numbers of both clinical and environmental samples (52 vs. 70, respectively), VNBII contained almost exclusively clinical isolates (60 vs. 3, respectively), which was a significant difference (p < 0.0001, Fisher’s exact test). Environmental isolates in both VNBI and VNBII were scattered across the phylogenetic tree and showed no evidence of clustering, although all three environmental VNBII isolates were relatively basal in the lineage (**Fig. 1**). SNP calling and phylogenetic analysis of 23 recently published Zambian isolates (Vanhove et al. 2016) showed a similar bias in environmental and clinical isolates in the two lineages; the Zambian mixed environmental and clinical VNB-A clade matched our VNBII lineage, while the VNB-B clade, which was entirely environmental samples, matched our VNBI lineage (**Supplemental Fig. S2**). Furthermore, VNBI had a greater proportion of *MAT***a** isolates than VNBII (33/122 vs. 9/64, respectively), although the difference was not significant.

Analysis of annotated whole-genome assemblies of representatives of the four lineages revealed very few gene gains or losses specific to VNBI and VNBII (**Supplemental Table S2**). Most gains and losses were hypothetical proteins, with the exceptions of a gain of an α-β hydrolase and loss of a phosphopyruvate hydratase and an alginate lyase in VNBI, and a gain of an L-iditol 2-dehydrogenase in VNBII. However, a large number of fixed polymorphisms were present between the two lineages, in addition to private unfixed alleles in each lineage and unfixed alleles shared between the two lineages (**Supplemental Fig. S3A**). Therefore small polymorphisms may underlie much of the phenotypic variation within and between these lineages.

To identify genomic features that might contribute to increased prevalence in clinical populations, we conducted a genome-wide association analysis (GWAS) of small polymorphisms and loss-of-function mutations across VNBI and VNBII. Correction for population stratification was included so that features that evolved at the base of VNBI would not overwhelm the analysis. The top 10 loci associated with the clinical phenotype included a number of known virulence factors and genes involved in oxidative stress response (**Table 2**). The top scoring locus was the intergenic region downstream of CNAG_07989 (p < 3.0×10^−7^, score test), which encodes a velvet family protein, and a deletion of that gene prevents filamentation (Chacko et al. 2015). Also identified were the intergenic region downstream of *LMP1,* a known virulence factor identified through laboratory passage of H99 (Janbon et al. 2014), *CCK2,* which encodes a casein kinase related to *CCK1* (essential for cell integrity and virulence (Wang et al. 2011)), and three genes, *PDX3, FOX21,* and *FHB1,* previously noted to respond to oxidative and nitrosative stress (de Jesús-Berríos et al. 2003; Upadhya et al. 2013).

**Table 2.** GWAS analysis reveals loci associated with the clinical phenotype in VNBI and VNBII. Two GWAS analyses were conducted. In the first, variants under 5% frequency were combined by gene or intergenic region (rare) while variants over 5% frequency were treated independently (common). In the second analysis, loss-of-function mutations were identified and combined by gene (LOF). Both analyses were conducted using GEMMA corrected for population stratification with a relatedness matrix (Zhou and Stephens 2012). The 10 most significant loci according to the score test across both analyses are shown.

| P value | Hit Type | Locus | Genes(s) |
| --- | --- | --- | --- |
| 2.02×10<sup>-7</sup> | LOF | CNAG_01348 | cyanate hydratase |
| 3.00×10<sup>-7</sup> | rare | intergenic: CNAG_07989-CNAG_07661 | velvet family protein, deletion prevents filamentation (Chacko et al. 2015); hypothetical protein |
| 3.22×10<sup>-7</sup> | common | intergenic: CNAG_07972-CNAG_06879 | chromosome 3 centromeric region |
| 1.19×10<sup>-6</sup> | rare | CNAG_05440 | pyridoxamine 5'-phosphate oxidase *PDX3*, downregulated under oxidative stress (Upadhyaya et al. 2013) |
| 2.44×10<sup>-6</sup> | rare | CNAG_03374 | DNA replication complex GINS protein *PSF1* |
| 2.91×10<sup>-6</sup> | common | intergenic: CNAG_06765-CNAG_06764 | low mating performance *LMP1*, deletion is avirulent (Janbon et al. 2014); short chain dehydrogenase *FOX21*, downregulated under oxidative stress (Upadhyaya et al. 2013) |
| 3.99×10<sup>-6</sup> | common | CNAG_06921 | hypothetical protein |
| 5.07×10<sup>-6</sup> | common | CNAG_05037 | hypothetical protein |
| 5.62×10<sup>-6</sup> | rare | CNAG_07427 | Casein Kinase *CCK2*, related to *CCK1* which is essential for virulence (Wang et al. 2011) |
| 6.00×10<sup>-6</sup> | rare | CNAG_01464 | Flavohemoglobin *FHB1*, nitrosative stress response (de Jesús-Berrios et al. 2003) |

We also examined geographical associations of the VNBI and VNBII lineages. VNBII isolates were isolated almost exclusively from patients treated at the Princess Marina Hospital in southern Botswana, while VNBI isolates were common in all four heavily sampled locations, including patients from both hospitals and the environment in south, northeast, and north-central Botswana (**Supplemental Table S3**). Further evaluation of the VNBI population revealed little evidence for genetic structure separating or connecting specific locations (**Supplemental Table S4**). We also examined the combined phylogeny (**Supplemental Fig. S2**) of the Botswana isolates with the previously sequenced Zambian isolates (Vanhove et al. 2016) to determine if these patterns extended outside of Botswana. Surprisingly, both VNBI and VNBII Zambian isolates largely clustered near the base of each lineage, suggesting some genetic separation between Botswanan and Zambian VNB lineages and that some geographic isolation may be occurring at the scale of country. The Zambian VNI isolates, on the other hand, were dispersed throughout our global VNIa and VNIb clades, though not the African VNIc clade, suggesting greater gene flow between VNI populations in the region.

### VNBI and VNBII differ in their ability to melanize and respond to oxidative stress

To determine if the distinct genetic backgrounds of VNBI and VNBII, or variants identified in the association analysis, correlated with any relevant phenotypes, we assayed 183 of the 186 sequenced VNBI and VNBII isolates for growth under a variety of stress conditions, along with a set of control isolates including the VNI reference strain H99. We specifically tested response to oxidative stressors H_2_O_2_ and paraquat, melanization capacity with L-DOPA, and resistance to the antifungal drug fluconazole (results in **Supplemental Table S5**). We did not identify any isolates with mutations in stress response genes *PDX3, FOX21,* or *FHB1* with extreme response phenotypes. However, we did identify significant phenotypic differences between VNBI and VNBII isolates and between clinical and environmental isolates. To control for the interaction of lineage and isolation source, we analyzed only clinical isolates for lineage comparisons and only VNBI isolates for isolation source comparisons. We also excluded all isolates that were completely unable to melanize in the comparison of melanization between environmental and clinical isolates to prevent any effect of sampling bias (Methods).

VNBI clinical isolates were significantly more melanized than VNBII clinical isolates (p < 0.00017, Mann-Whitney *U* test, **Fig. 4A**), and significantly more resistant to both H_2_O_2_ (p < 0.0049, Mann-Whitney *U* test, **Supplemental Fig. S4A**) and paraquat (p < 0.042, Mann-Whitney *U* test, **Supplemental Fig. S4B**), but not fluconazole (p < 0.78, Mann-Whitney *U* test, **Supplemental Fig. S4C**). Within VNBI, environmental isolates were more heavily melanized (p < 0.00047, Mann-Whitney *U* test) than clinical isolates and more resistant to both H_2_O_2_ (p < 0.00098, Mann-Whitney *U* test) and paraquat (p < 4.1×10^−5^, Mann-Whitney *U* test). The groups did not significantly differ in their resistance to fluconazole (p < 0.071, Mann-Whitney *U* test).

**Figure 4.**
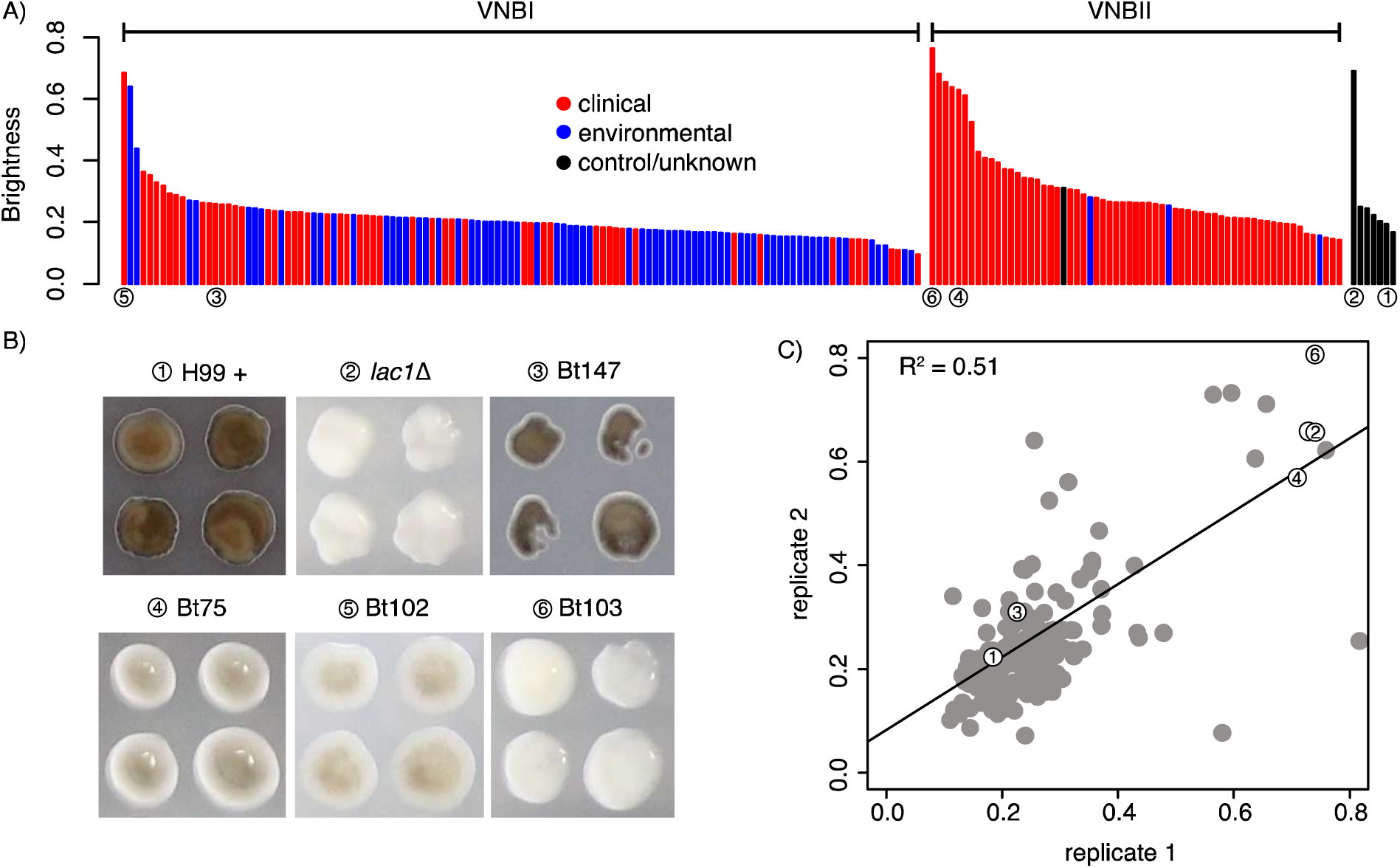
Phenotypic and GWAS analyses demonstrate that VNBI isolates have a greater ability to melanize than VNBII isolates and identify *BZP4*-deficient isolates as having reduced melanization capacity. A) Isolates were provided with L-DOPA and colony brightness was assayed; isolates with the lowest brightness are the most melanized. Clinical isolates are shown in red, environmental isolates are shown in blue, control/unknown isolates are shown in black, and isolates described in (B) are underscored with numbered circles. VNBI clinical isolates were significantly more melanized than VNBII clinical isolates (p < 0.00017) and VNBI environmental isolates were significantly more melanized than VNBI clinical isolates (p < 0.00047). For the latter comparison, the least melanized (brightness >= 0.6) samples were excluded to prevent an effect of sampling bias. B) GWAS analysis identified loss-of-function mutations in *BZP4* as being associated with a lack of melanization. The four isolates with *BZP4* loss-of-function mutations are shown here in the L-DOPA assay, along with the positive control H99 and the negative control *lac1∆*. C) A strong correlation of melanization was present between replicates. Isolates shown in (B) are indicated with numbered circles.

We then conducted a second GWAS analysis to identify genotypic variants associated with increased or decreased responses in the phenotypic assays (**Supplemental Tables S6-S9**). These analyses revealed that loss-of-function mutations in *BZP4* (CNAG_03346) were highly associated with reduced melanization ability (p < 3.43×10^−9^, score test). Four clinical isolates, two from VNBI and two from VNBII, each had a different loss-of-function mutation in *BZP4.* Three of these isolates had a reduced melanization ability similar to that of a *lac1* deletion mutant, the primary gene required for melanization (Salas et al. 1996), while the fourth isolate showed a slightly reduced melanization potential (**Fig. 4A-C**). *BZP4* was previously associated with a reduced melanization capacity in a deletion screen of *C. neoformans* transcription factors (Jung et al. 2015). We then further screened genes previously categorized as having melanization defects in another deletion screen (Liu et al. 2008) and identified a second gene, *CHO2* (CNAG_03139) at lower rank in the association analysis (p < 0.0083, score test). Loss-of-function of this gene is present in all VNBII isolates and could be related to the reduced melanization capability of VNBII. While GWAS analysis did not produce any promising candidates for fluconazole resistance, copy number analysis revealed a duplication of drug target *ERG11* in the four most resistant isolates (Bt65, Bt89, MW-RSA3179, and MW-RSA2967).

### Population genomic analysis identifies extensive loss of diversity in VNBI

To better understand genetic diversity at the lineage level, we calculated a number of population genetic statistics for VNI, VNBI, and VNBII, including π, Tajima’s D, and F_ST_ (**Table 1**). Genome-wide calculations of the diversity metric π revealed that VNBI and VNBII had nearly twice the genetic diversity of VNI (**Table 1**). However, when diversity was plotted along each chromosome, VNBI and VNBII appeared quite different. As shown in **Fig. 5A** for chromosome 5, the VNI group contained long tracts of low diversity while VNBII contained predominantly high diversity regions, as expected. By contrast, VNBI showed a unique pattern with high diversity regions interspersed with long tracts of low diversity. These regions of low diversity correspond with tracts of low Tajima’s D, which suggests an excess of rare alleles resulting from a population bottleneck or recent selective sweep. Confirming this, both VNI and VNBI had significantly more rare alleles across the entire genome (<2% frequency) than VNBII (p<2.2x10^−16^ for both comparisons, Fisher’s exact test, **Supplemental Fig. S3B**). To determine if this pattern appeared in all chromosomes, we plotted the distribution of Tajima’s D values for each chromosome individually (**Fig. 5B**). While VNI and VNBII have unimodal distributions for most chromosomes, VNBI has a bimodal distribution of Tajima’s D across nearly all chromosomes, with the possible exception of chromosome 6. However, when we evaluated genes located in the regions of reduced Tajima’s D we found no significant functional enrichment in any of the lineages.

**Figure 5.**
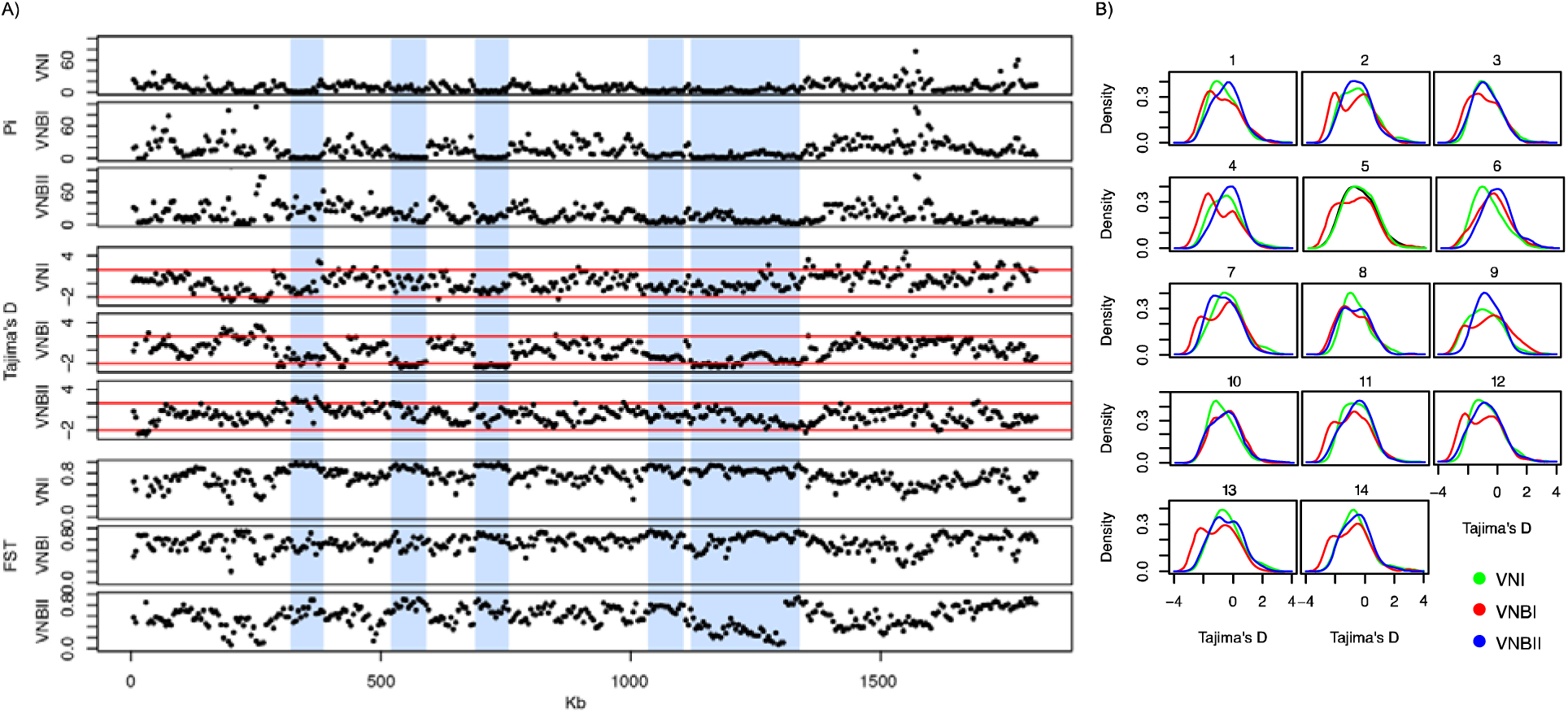
Population genomic analysis revealed long tracts of low genetic diversity and Tajima’s D in VNI and VNBI. A) Low diversity tracts are shown here in chromosome 5. In VNBI, these tracts are interspersed between regions of high diversity. Vertical blue bars highlight these areas of reduced diversity and Tajima’s D in VNBI, which are generally accompanied by high F_ST_ between populations. Statistics shown include π, Tajima’s D, and F_ST_. B) Density distribution of Tajima’s D across all 14 chromosomes for VNI, VNBI, and VNBII. VNI and VNBII show predominantly unimodal distributions for most chromosomes, while VNBI shows a bimodal distribution for all chromosomes except 6.

In most cases, when two lineages had regions of low diversity and Tajima’s D, they also had high F_ST_. However, there was one notable exception to this on chromosome 5, where a region of reduced diversity in both VNBI and VNBII included a region of high F_ST_ abruptly adjacent to a region of low F_ST_ (**Supplemental Fig. S5**). The low F_ST_ region includes CNAG_07442, which encodes a phosphatidylglycerol transfer protein that was the most abundant antigen recovered from an experimental mouse vaccine using strain cap59 (Specht et al. 2015), suggesting a similar antigen would be found in both lineages. The high F_ST_ region included CNAG_01058, a gene involved in capsule maintenance (Brown et al. 2014) and CNAG_01060, a potential substrate of the *FBP1* gene required for proliferation in macrophages and dissemination (Liu and Xue 2014).

### Inositol and other sugar transporters are targets of selection in all VN lineages

As Tajima’s D can identify effects of both selection and demography (Tajima 1989), we used the composite likelihood ratio test (CLR) (Nielsen et al. 2005), which is specific to selection, in a genomic scan for regions under selection in each lineage (**Supplemental Fig. S6**). Surprisingly, there was very little overlap between the genes in regions under positive selection and the genes in regions of reduced diversity in any of the lineages, suggesting that selective sweeps were not fixing genes under selection. To further evaluate genes under positive selection, we identified genes in regions with the top 5% of CLR values for each of VNI, VNBI and VNBII. A functional enrichment analysis of these genes revealed that major facilitator superfamily (MFS) transporters, and in particular the sugar transporter subset, were under selection in all lineages (**Supplemental Table S10**). Overlaying these genes onto a phylogeny of all sugar transporters in the reference strain H99 showed the selection was concentrated in transporters of inositol, xylose, maltose, α-glucosides, lactose, and galactose (**Supplemental Fig. S7**). Six inositol transporters were found to be under selection in at least one lineage, including *ITR1A* (CNAG_04552) which was under selection in all three lineages (**Table 3**). Furthermore, genes involved in inositol catabolism, including four copies of inositol 2-dehydrogenase and one copy of inositol oxygenase were also under selection, notably in VNI where all five genes were under selection. Other groups of genes under selection with enriched functions include monoxygenases in VNI and β-glucosidases in VNBII.

**Table 3.** Inositol transport and utilization genes under selection in individual lineages. Genes were present in the top 5% of windows evaluated with the CLR test for each lineage.

| H99 Locus | Gene | VNI | VNBI | VNBII |
| --- | --- | --- | --- | --- |
| CNAG_00097 | inositol transporter *ITR1* | + | - | + |
| CNAG_04552 | inositol transporter *ITR1A* | + | + | + |
| CNAG_00864 | inositol transporter *ITR2* | - | + | - |
| CNAG_05377 | inositol transporter *ITR3* | + | - | - |
| CNAG_00867 | inositol transporter *ITR3A* | - | + | - |
| CNAG_05667 | inositol transporter *ITR13B* | - | + | - |
| CNAG_05329 | *myo*-inositol 2-dehydrogenase | + | - | + |
| CNAG_05320 | inositol 2-dehydrogenase | + | + | - |
| CNAG_05321 | inositol 2-dehydrogenase | + | + | - |
| CNAG_06530 | inositol 2-dehydrogenase | + | - | - |
| CNAG_05316 | inositol oxygenase | + | - | - |

Subtelomeric regions of *Cryptococcus* have been previously highlighted as regions of high diversity heavily enriched for transporters (Chow et al. 2012). To determine if subtelomeric regions showed signatures of selection, we searched for enrichment of windows under selection in the outer 40 kb of each chromosome for VNI, VNBI, and VNBII. Chromosomes 1,4, 5, and 14 showed subtelomeric enrichment of selection in all three examined lineages, and all chromosomes except 2 and 6 showed enrichment in at least one lineage (**Supplemental Table S11**). This suggests subtelomeric regions encode numerous targets of selection across the VNI, VNBI, and VNBII lineages. Enrichment analysis comparing genes under selection in the subtelomeric regions to the remainder of the genome showed the same patterns as the genome-wide selection analysis. Additionally, an enrichment of fungal transcription factors was identified (**Supplemental Table S12**), including *FZC20, FZC41, SIP402,* and *FZC22,* the last of which has been shown to have reduced virulence in mice when deleted (Jung et al. 2015).

### Mating type loci *MAT*α and *MAT*a have distinct evolutionary trajectories

The mating type locus also showed strong signatures of selection. To better understand the evolution of the *MAT* locus, we re-called SNPs against complete assemblies of each of the two highly diverged mating types (*MATα* and *MAT****a***) (Lengeler et al. 2002) and inferred phylogenies of each mating type separately (**Fig. 6**, Methods). The *MATα* phylogeny largely matched the whole-genome phylogeny, with one major exception: VNBI *MATα* appeared paraphyletic with respect to VNBII (**Fig. 6A, B**), with two distinct alleles—one comprised of some VNBI and the other clade comprised of the remaining VNBI plus all VNBII (bootstrap support: 100%). These distinct clades did not map to specific sub-clades of VNBI in the whole-genome phylogeny but rather were intermixed.

**Figure 6.**
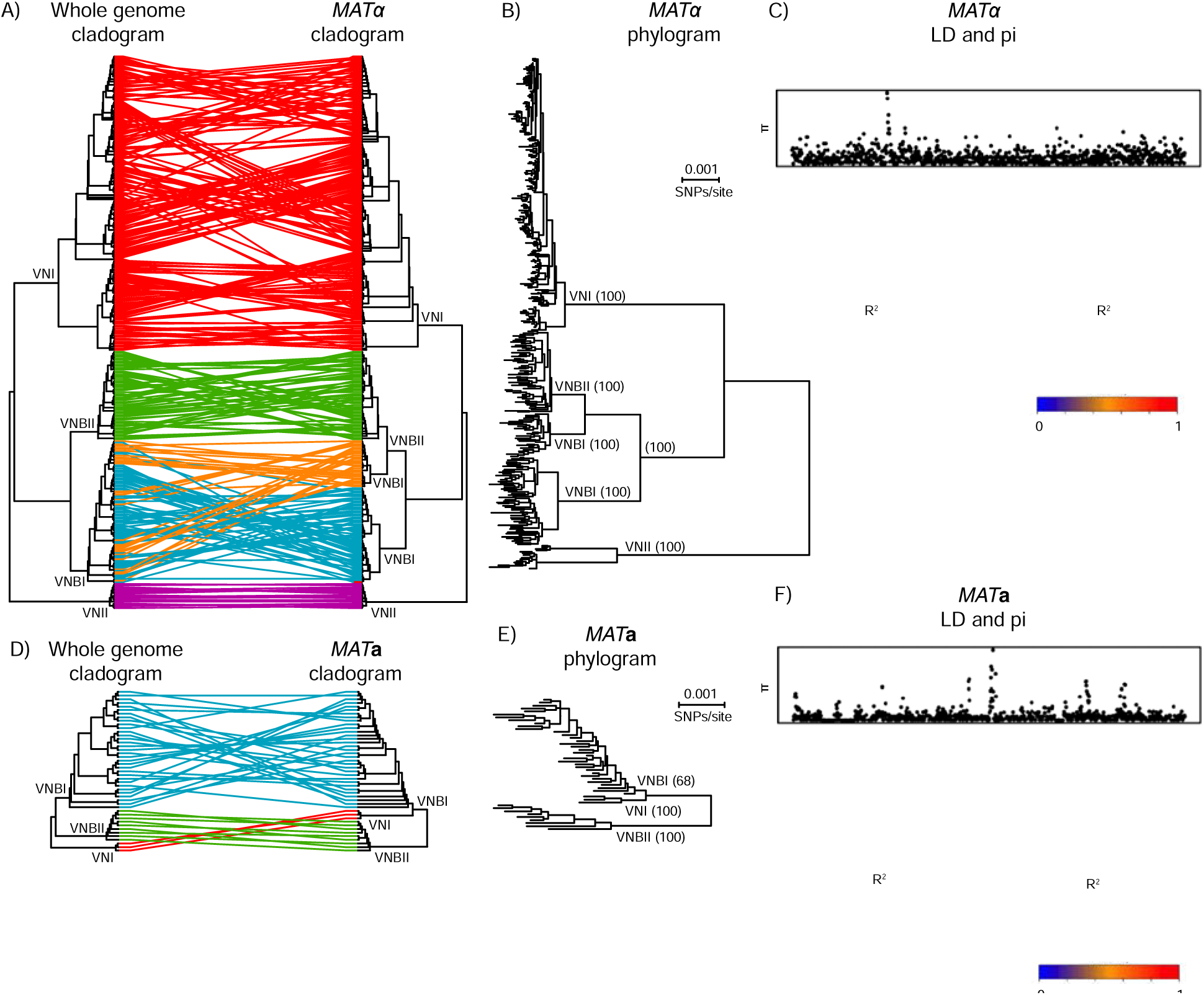
Phylogenetic and linkage analyses reveal distinct evolutionary trajectories of the *MAT* locus alleles. A) The topology of the whole-genome phylogeny and *MATα* phylogeny are compared as cladograms. VNBI isolates contain two distinct a alleles; B) *MATα* phylogram showing branch lengths and bootstrap support for major clades; C) Linkage disequilibrium (R^2^) and diversity (π) across the *MATα* locus; D) Topology of the whole-genome phylogeny and *MAT***a** phylogeny are compared as cladograms; E) *MAT***a** phylogram showing branch lengths and bootstrap support for major clades. VNI is close sister-group to VNBI, in contrast to the whole-genome phylogeny, and there is limited support for monophyly of the VNI *MAT***a** allele with respect to VNBI (bootstrap support: 68%); F) Linkage disequilibrium (R^2^) and diversity (π) across the *MAT***a** locus.

The *MAT***a** phylogeny also differed from the whole-genome phylogeny (**Fig. 6D, E**) in that VNI and VNBI were closely related sister groups and separated by a large distance from VNBII (bootstrap support: 100%). Bootstrap support for the monophyly of VNBI with respect to VNI was limited (68%), indicating the possibility that the VNI *MAT***a** allele was derived from a basal VNBI allele. Additionally, alleles co-segregating VNBI *MAT***a** isolates with VNI and VNBII *MAT***a** isolates extend about 5.3 and 3.5 kb, respectively, upstream of the 5’ end of the *MAT* locus (**Supplemental Fig. S8**). This suggests a local introgression of the *MAT***a** locus from VNBI into VNI, consistent with previous reports of hotspots that flank the *MAT* region (Hsueh et al. 2006; Sun et al. 2012).

The *MATα* and *MAT***a** alleles also showed distinct patterns of genetic diversity. Both alleles showed spikes of high genetic variation in a small number of hotspots (**Fig. 6C, F: π**); all of the regions corresponded to long terminal repeats in the annotations of the mating type loci, suggesting only limited variation has accumulated in functional genes. Within *MATα*, VNI alleles had significantly fewer singleton polymorphisms than either VNBI or VNBII, resulting in shorter terminal branch lengths (**Fig. 6B**; Wilcoxon rank sum test, p < 3.8×10^−5^ and p < 5.2×10^−6^, respectively). While this could be the result of mixing within VNI but not VNBI or VNBII, the same pattern was observed in the whole-genome data (**Fig. 1**; Wilcoxon rank sum test, p < 2.2×10^−16^ and p < 1.7×10^−8^), suggesting an overall increase in genetic variation in VNBI or VNBII relative to VNI may be responsible for the difference. Between *MATα* and *MAT***a***, MAT***a** alleles had significantly more singleton polymorphisms than *MATα*, even when VNI isolates were excluded (**Fig. 6B, E**; Wilcoxon rank sum test, p < 1.3×10^−6^), which could indicate less mixing between *MATα* alleles. Extensive linkage disequilibrium was seen in both *MAT* alleles (**Fig. 6C, F**: R^2^), suggesting that despite recombination occurring at the whole-genome level within VNI, VNBI, and VNBII, limited or no recombination was occurring within the mating type loci. In total, these analyses suggest that each mating type locus in *C. neoformans* has followed a distinct evolutionary trajectory from the remainder of the nuclear genome.

## Discussion

Phylogenomic analysis presented here provided strong support for a deep split separating the VNB lineage in VNBI and VNBII, and further population genomic analysis revealed a lack of recombination between the two. These lineages also differed in their ability to be recovered from the environment, respond to oxidative stress, and melanize, and to a lesser extent the presence of both mating types (*MAT***a** was less common in VNBII). Based on these genomic and phenotypic differences, we propose that VNBI and VNBII should be considered two separate lineages. The lower frequency of isolates from environmental sources in VNBII could be due to an increased ability for VNBII isolates to colonize humans; there also may be geographic or niche differences in VNBII environmental populations. Despite the fact that melanization is used to identify isolates from the environment (*i.e.,* light brown isolates are detected but pure white isolates are not), it is unlikely that decreased melanization of VNBII caused the group to be under-sampled as most VNBII isolates melanized to at least some degree. Further genotypic and phenotypic evaluation will be required to better delineate and understand the clinical ramifications of the differences between VNBI and VNBII.

Both “into” and “out of” Africa hypotheses have been proposed for VNI (Litvintseva et al. 2011). However, given the existence of the sub-Saharan Africa endemic VNIc and the presence of African isolates spread across VNIa and VNIb, the “into” hypothesis seems unlikely. More likely, the three VNI sub-lineages diversified and expanded within Africa and then two were globally disseminated at a later date, possibly by migration of birds and humans. Why VNI but not VNBI or VNBII would be globally dispersed is unclear. While the data presented here support an African origin for VNB, a small number of South America VNB isolates have been described, including a previous analysis of nuclear loci that placed two of these isolates close to Bt22 (Ngamskulrungroj et al. 2009), identified here as a VNBI isolate. Comprehensive sequencing of South American VNB isolates will be required to confidently assess the origins of VNB diversity.

Given the lack of overlap between the tracts of low Tajima’s D and the regions identified as under selection, we hypothesize that the loss of diversity in VNI and VNBI are the result of previous demographic events, such as population bottlenecks, rather than selective sweeps fixing genes under selection. The global VNI lineage, with low overall diversity, likely emerged from a small founder population that may have undergone repeated bottlenecks while spreading geographically. Why the VNBI lineage, endemic to southeastern Africa, would have undergone a major demographic event, is unclear, as it appears to be the most well-represented lineage in current environmental sampling.

Previous analyses have highlighted the presence of recombination between isolates of different lineages based on MLST gene analysis (Litvintseva et al. 2006; Chen et al. 2015). The resolution possible with whole genome data demonstrates that many of these isolates predicted to be recombinant by analysis of single genes such as *SOD1* are predominantly a single ancestry based on whole genome analyses (*e.g.,* Bt65, Bt109). The detection of small introgressions in VNBI, VNBII, and African VNI isolates suggests that a limited amount of recombination is occurring in Africa where the three groups overlap in distribution. Given the clinical significance of genes in these regions such as *ERG11,* better understanding of the phylogenetic and geographic distribution of introgressions may be of clinical importance.

VNI showed a level of intragroup recombination comparable to VNBI and VNBII. This was unexpected as both mating types are common in the VNB lineages, suggesting sexual reproduction would dominate in VNB, while the **a** mating type is rare in VNI, suggesting predominance of asexual reproduction in VNI. However, the high level of recombination seen in VNI may indicate unisexual mating in VNI is nearly as common as bisexual mating in VNBI and VNBII. It has been shown in *C. neoformans* var. *neoformans* (serotype D) isolates that recombination is similar in unisexual and bisexual mating (Sun et al. 2014; Fu et al. 2015).

These population genomic analyses can also be used to evaluate lineage and species boundaries that are currently under debate for *Cryptococcus* (Hagen et al. 2015; Kwon-Chung et al. 2017). A multi-locus viewpoint is critical for a comprehensive understanding of recombination and incomplete lineage sorting within a species, as recently shown for *Alternaria* (Stewart et al. 2014). Despite high bootstrap support for the monophyly of the four population subdivisions (VNI, VNII, VNBI and VNBII) in the whole-genome phylogenetic tree, the detection of recombination between the lineages, including small introgressions, suggests that isolates from the different lineages mate and form recombinant progeny; further analysis including of hybrid strains (Chen et al. 2015) will help clarify if these lineages represent varieties or sibling species. While whole-genome analyses may largely recapitulate the phylogenetic relationships revealed from MLST studies, they can also reveal complex genetic exchange within a species at a level of detail not possible in smaller scale studies.

Phylogenetic analyses of the *MAT* alleles suggest that these alleles have followed distinct evolutionary trajectories from the remainder of the nuclear genome, involving incomplete sorting or genetic mixing following divergence of the groups, as seen in the paraphyly of the VNBI MATα allele and close relationship of the VNI *MAT***a** allele to VNBI. These observations suggest an outline of the evolution of mating in these groups: the presence of both mating types is ancestral in VNBI and VNBII, and sexual reproduction in these groups involves opposite as well as potentially same sex mating. However, VNI may have diverged with only the MATα mating type, possibly due to a small founding population size. This would have provided evolutionary pressure for unisexual reproduction, as is suggested by the evidence of extensive recombination within the nuclear genomes of VNI isolates. Following the divergence of VNI from VNBI and VNBII, we speculate that VNI then re-acquired the *MAT***a** allele from a VNBI-like ancestor.

The selection and association analyses, along with the phenotypic assays, provided clues to both selection pressures experienced by *C. neoformans* in the environment and which of these traits may translate to pathogenicity in humans. Selection on both inositol transport and utilization could affect the lineage-specific host range of *C. neoformans* and easily translate to altered virulence levels in humans, as inositol is abundant in the human brain (Fisher et al. 2002) and is required by *C. neoformans* for virulence (Shea et al. 2006) and mating (Xue et al. 2010). Gene expansions of inositol transporters in *C. neoformans* have been previously proposed to have evolved for adaptation growth on trees yet preadapted *C. neoformans* for growth in the human brain (Xue et al. 2010; Xue 2012). Selection on xylose facilitators suggests potential adaptation to different tree hosts with different chemical compositions. GWAS analysis also revealed a gene known to be involved in the yeast-hyphal transition. As the inability to maintain yeast phase is known to correlate with decreased virulence (Wang et al. 2012), it is easy to hypothesize that natural adaptations to environments where increased time in the yeast state would be selected for could result in isolates with increased virulence in humans. Future genomic and phenotypic analyses focused on VNI may help better understand the specific properties that enabled the global dispersal of this lineage.

It was initially surprising that within VNBI, environmental isolates were more resistant to oxidative stress and more heavily melanized than clinical isolates, particularly given that melanization is a known virulence factor in humans. However, the diversity of stressors in the environment is likely greater than within the human host, and it has already been hypothesized that the ability of *C. neoformans* to survive in macrophages was derived from an ability to defend against predatory protozoa in the environment (Steenbergen et al. 2001). Further experiments are needed to disentangle the stresses faced in these two contexts and the differential ability of environmental and clinical isolates to respond to these stresses. Here, GWAS analysis linked loss-of-function mutations in the transcription factor *BZP4* to significantly reduced melanization in a number of unrelated clinical isolates, highlighting the potential clinical impact of this complex interplay. In sum, these data provide insight into how selection pressure over the evolutionary history of *C. neoformans* may have pre-adapted to successful invasion of the human nervous system, and suggests many pathways to explore for differential virulence within standing genetic diversity or outbreaks.

## Methods

### Sample Selection

We selected 387 isolates for analysis: 185 from VNI, 186 from VNB, and 16 from VNII. We sought to represent isolation sources and mating types as evenly as possible for each of VNI and VNB, and therefore preferentially chose isolates containing *MAT***a** and those isolated from environmental sources, as both are less frequently collected, particularly for VNI. We also included a limited number of VNII isolates, but did not focus on this lineage as it is rarely isolated. The majority of isolates, particularly VNB isolates, were collected in Botswana, and more specific collection details of these isolates are given in **Supplemental Material** and a previous MLST analysis (Chen et al. 2015).

### Sample Preparation, Sequencing, and Variant Identification

Samples were prepared for DNA extraction as described (**Supplemental Material**). Genomic DNA was sheared to ~250 bp using a Covaris LE instrument and adapted for Illumina sequencing as previously described (Fisher et al. 2011). Libraries were sequenced on an Illumina HiSeq to generate 101 base reads. Reads were aligned to the *C. neoformans* var. *grubii* H99 assembly (Genbank accession GCA_000149245.2) using BWA-MEM version 0.7.12 (Li 2013). Variants were then identified using GATK version 3.4 (McKenna et al. 2010) as described (**Supplemental Material**), and functionally annotated with SnpEff version 4.2 (Cingolani et al. 2012). Read data from 46 previously sequenced Zambian isolates (Vanhove et al. 2016) was downloaded from NCBI (BioProject PRJEB13814) and variants were called for all samples as described above.

### Phylogenetic Analysis

For phylogenetic analysis, the 1,069,080 sites with an unambiguous SNP in at least one isolate and with ambiguity in at most 10% of isolates were concatenated; insertions or deletions at these sites were treated as ambiguous to maintain the alignment. Phylogenetic trees were estimated using RAxML version 8.2.4 (Stamatakis 2014) under the GTRCAT model in rapid bootstrapping mode. We then placed the 46 previously sequenced Zambian isolates (Vanhove et al. 2016) in phylogenetic context with the isolates in this study using FastTree version 2.1.8 compiled for double precision (Price et al. 2010).

### Population Genomic Analysis

Population structure was investigated using multiple approaches. Principal components analysis was run on all variants using SMARTPCA (Patterson et al. 2006). Major ancestry subdivisions were delineated using STRUCTURE (Pritchard et al. 2000) v2.3.4 in the site-by-site mode and k=4, with 15% of the positions randomly subsampled from those containing at least 2 variants and missing in fewer than 5% of isolates. Identification of VNI subdivisions and smaller inter-lineage introgressions using STRUCTURE were performed as described (**Supplemental Material**). Population genomic metrics were calculated using vcftools version 1.14 (Danecek et al. 2011) and PopGenome (Pfeifer et al. 2014); further details are given in **Supplemental Material**. The CLR selection metric was also calculated using PopGenome, and enrichment tests were conducted using Fisher’s exact test corrected with the Benjamini-Hochberg method for multiple comparisons (**Supplemental Material**).

### Association Analysis

For each phenotype, two matrices of these features were constructed from the variant calls of all isolates. In the first matrix, rare variants (those at ≤ 5% frequency) were collapsed by gene or intergenic region, while common variants (those at > 5% frequency) were considered independently. In the second matrix, we assessed whether each gene had a loss-of-function mutation (defined as s “HIGH” impact mutation by SnpEff), regardless of the frequency of the mutation. Each variant matrix was then analyzed with GEMMA (Zhou and Stephens 2012) using a univariate linear mixed model and a relatedness matrix to account for population stratification. Score test results of both analyses were then combined and examined. *ERG11* and *AFR1* copy number of drug resistant isolates were estimated as the normalized read depth of these genes from whole genome BWA-MEM alignments, where the per base depth calculated using samtools (Li et al. 2009) v1.3 mpileup was summed across each gene and normalized to the average genome depth.

### Phenotypic Assays

Freezer stocks were grown, plates, and pinned in 1536 array format as described (**Supplemental Material**); each single colony selected from the grown freezer stock became a 4 x 4 block of 16 colonies in two 1536 arrays. Four conditions were assayed: 5.6 mM H_2_O_2_, 1.0 mM paraquat, 10 μg/mL fluconazole, and 0.1 mg/mL L-DOPA. Arrays were incubated at 30°C and images were acquired one-, two-,and three-days post pinning. Day two images were analyzed using gitter (Wagih and Parts 2014) to assess either colony size in the case of H_2_O_2_, paraquat of fluconazole, or brightness in the case of L-DOPA. Each condition was tested in either one (H_2_O_2_), two (L-DOPA and fluconazole) or three (paraquat) independent experiments. All independent experiments replicated well (R^2^, L-DOPA = 0.51, fluconazole = 0.59, paraquat = 0.47-0.63). We then identified differences between VNBI and VNBII clinical isolates, and VNBI environmental and clinical isolates, using the Mann-Whitney *U* test. As *C. neoformans* is identified in the environment (but not in the clinic) by the ability to melanize to some degree *(i.e.,* light brown isolates are detected but pure white isolates are not), in the comparison of melanization of environmental and clinical VNBI isolates, we excluded isolates that were unable to melanize (brightness >= 0.6) because the sampling of those isolates would be biased towards clinical samples.

### Mating Type Analysis

Analyses of the mating type loci were done with the same methodology as the whole genome analyses with the following two exceptions. Reads were mapped to the appropriate high quality locus-specific reference, either the H99 *MAT*α allele (Genbank accession AF542529) or the 125.91 *MAT***a** allele (Genbank accession AF542528). Second, linkage disequilibrium was calculated on the site level rather than in windows, and visualized using the R package LDheatmap (Shin et al. 2006).

### Data Access

Genbank accession numbers of all sequencing data generated for this study are listed in **Supplemental Table S1**.

## Acknowledgments

We would like to thank Jose Munoz for providing helpful comments on the manuscript. This project has been funded in whole or in part with Federal funds from the National Institute of Allergy and Infectious Diseases, National Institutes of Health, Department of Health and Human Services, under Grant Number U19AI110818 to the Broad Institute.

Support to J.R.P. came from Public Health Service Grants AI73896, AI93257. The use of product names in this manuscript does not imply their endorsement by the US Department of Health and Human Services. The finding and conclusions in this article are those of the authors and do not necessarily represent the views of the CDC.

## Author Contributions

C.A.C., A.L., and J.P. conceived and designed the project. M.H. provided the clinical isolates. C.G., T.Y., Y.C., and A.M.J. performed the laboratory experiments. C.A.D., S.M.S., C.-H.Y., C.A.C., S.S., J.H., and C.G. analyzed the data. C.A.D., C.A.C., and A.L. wrote the paper. C.A.C., J.P., and J.L.T. supervised and coordinated the project.

## Ethics Statement

The clinical isolates were approved under Pro00029982 from the Duke Institutional Review Board in USA and the Review Boards of Nyangabgwe and Princess Marina Hospitals of Botswana.

